# Quercetin ameliorates mitochondrial dysfunction and mitigates methamphetamine-induced anxiety-like behavior

**DOI:** 10.1101/2021.06.29.450268

**Authors:** Fengrong Chen, Jiaxue Sun, Yongjin Zhang, Yicong Dai, Zherui Zhang, Cheng Chen, Lei Zou, Zunyue Zhang, Hongjin Wu, Weiwei Tian, Yu Xu, Huayou Luo, Juehua Yu, Kunhua Wang

**Author notes:** Corresponding authors: Kunhua Wang and Juehua Yu NHC Key Laboratory of Drug Addiction Medicine, The First Affiliated Hospital of Kunming Medical University, Kunming, 650032, Yunnan, China.

## Abstract

Methamphetamine (MA) abuse results in neurotoxic outcomes, including increased anxiety and depression, during both MA use and withdrawal. Although numerous studies have reported an association between MA exposure and anxiety, the underlying mechanism remains elusive. In this study, escalating dose of MA was used to establish an MA-treated mouse model presenting anxiety behavior. RNA seq was then performed to profile the gene expression patterns in the hippocampus (HIPP). Differentially expressed genes (DEGs) were identified and function enrichment analysis was conducted to explore the underlying mechanisms. Quercetin as an mitochondria protector was used *in vivo* and *in vitro*. The C57BL/6J mice were co-treated with 50 mg/kg Quercetin and escalating MA. Anxiety behavior was evaluated by utilizing the elevated plus maze and the open field test. Transmission electron microscopy and immunohistochemistry were conducted to study the pathology of MA-inducced anxiety. The effects of MA and Quercetin on astrocytes were investigated by fluorescence staining, transmission electron microscopy, flow cytometry, and oxygen consumption rate. Western blot and qPCR were performed to analyze altered protein and gene levels of HIPP in mice and astrtocytes. The results demonstrated that forteen upregulated differentially expressed genes were identified and significantly enriched in signaling pathways related to psychiatric disorders and mitochondrial function. Interestingly, we found that quercetin was able to alleviate MA-induced anxiety-like behavior by improving neuron number and mitochondria injury. Mechanistically, quercetin can mitigate aberrant mitochondrial morphology and mitochondrial dysfunction not only by decreasing the levels of total cytoplasmic reactive oxygen species (ROS), mitochondria-derived ROS (mtROS), and mitochondrial membrane potential (MMP), but also increasing the oxygen consumption rate (OCR) and mitochondrial ATP production *in vitro*, indicating Quercetin ameliorated MA-induced anxiety-like behavior by modifying mitochondrial morphology and function. Furthermore, quercetin reversed OPA1 and DRP1 expression in astrocytes, and mitigated astrocyte activation and the release of inflammatory factors, which can trigger neuronal apoptosis and synaptic loss. Taken together, we provided evidence showing that MA can induce anxiety-like behavior via the induction of oxidative stress and mitochondrial dysfunction. Quercetin exerted antipsychotic activity through mitochondrial modulation, suggesting its potential for further therapeutic development in MA-induced anxiety.

## INTRODUCTION

As a highly addictive psychostimulant drug, methamphetamine (MA) abuse is an increasingly common worldwide phenomenon, resulting in significant physical, behavioral, cognitive, and psychiatric outcomes (Homer et al., 2008; Meredith et al., 2005; Glasner-Edwards et al., 2014). Epidemiological studies have shown that amphetamine-type stimulants represent the most widely used illicit drugs in the world after cannabis, with ≤51 million global users between the ages of 15 and 64 years (Wearne et al., 2018). Among abusers, 72%–100% experience MA-induced psychotic reactions (Srisurapanont et al., 2003; Smith et al., 2009), and 30.2% of chronic MA users are diagnosed with anxiety (Hellem et al., 2016).

Human neuroimaging studies have shown that MA users experience significant changes in multiple brain regions, including the orbitofrontal cortex, striatum, amygdala, hippocampus, and insula, which are involved in a variety of functional networks, including the salience network, limbic system, and frontostriatal circuit (May et al., 2020; Uhlmann et al., 2016). Evidence has demonstrated that MA administration can evoke changes in behavior, synaptic transmission, and volume in the hippocampus and cortex (Golsorkhdan et al., 2020; Uhlmann et al., 2016), resulting in the dysregulation of neurotransmitters and their receptors in these regions (Uhlmann et al., 2016), often accompanied by oxidative stress, apoptosis, and autophagy (Huang et al., 2017).

A limited number of studies have preliminarily demonstrated that both oxidative stress and mitochondrial dysfunction may play roles in the pathology of anxiety (Chang et al., 2007; Kohno et al., 2018). Excessive dopamine induced by MA is thought to trigger the overproduction of reactive oxygen species (ROS) by the mitochondria and relevant enzymes, exacerbating neurodegenerative diseases (Shin et al., 2017), suggesting that mitochondrial functional processes may play major roles in the brain abnormalities associated with MA-induced anxiety. Increasing experimental evidence has supported the existence of a link between mitochondrial dysfunction, brain dysfunction, and neuropsychiatric disorders (Pei et al., 2018; Wallace et al., 2017; Filiou et al., 2019). Suboptimal mitochondrial function would be vulnerable to the stress-associated depletion of the brain’s energy resources, resulting in the development of psychiatric disorders (e.g., anxiety and depression) (Morava et al., 2013). However, the mechanisms underlying the etiology of MA-induced anxiety remain poorly understood.

Research has suggested that the treatment of co-occurring psychiatric disorders, including depression and anxiety, may also be important for preventing relapses (Su et al., 2017; Glasner-Edwards et al., 2010). A substantial amount of literature has demonstrated the efficacy of both first- and second-generation antipsychotic drugs for the treatment of psychotic symptoms associated with MA-induced depression and anxiety (Chiang et al., 2019; Shoptaw et al., 2009; Wang et al., 2016). However, adverse events are frequently reported in these studies. Quercetin, which is a flavonoid-type secondary metabolite found in foods and medicinal plants, is presumed to have antioxidant, anti-inflammatory, immunoprotective, and anti-carcinogenic effects and was shown to mitigate anxiety-like behaviors in mice by modulating oxidative stress and monoamine oxidase activity (Dhiman et al., 2019), the preventing antioxidant enzyme impairment, regulating serotonergic and cholinergic neurotransmission, and decreasing neuroinflammation and neuronal apoptosis (Samad et al., 2018; Kosari-Nasab et al., 2019). Quercetin has been confirmed to be safe when used as a single compound in dietary supplements in both animal and human studies, and adverse effects following supplemental quercetin intake have rarely been reported (Andres et al., 2018). Despite evidence that quercetin serves as an oxidative stress and inflammatory modulator, no research has examined the effects of quercetin on anxiety-like behaviors induced by chronic MA.

In this study, we aimed to determine whether quercetin intervention following chronic MA use can mitigate anxiety-like behaviors by exploring the underlying molecular mechanisms and targeted brain regions, combined with RNA-sequencing (RNA-seq). We first constructed an animal model of MA-induced anxiety. The HIPP were isolated, and RNA-seq was performed to identify differentially expressed genes (DEGs) between the MA treated and control groups. Subsequently, the potential underlying pathways were analyzed by Kyoto Encyclopedia of Genes and Genomes (KEGG) analyses, and the identified DEGs were validated by quantitative reverse transcription-polymerase chain reaction (qPCR). Finally, in view of the functional enrichment, we assessed the antianxiety effects of quercetin in MA-treated mice to explore the underlying mechanism. These findings will contribute to a better understanding of the molecular mechanisms of anxiety induced by MA and allow for the therapeutic potential of quercetin against MA-induced anxiety to be assessed.

## MATERIALS AND METHODS

### Animals and treatment

Mice were housed with a 12-h light/dark cycle (lights on at 7:00 AM). Behavioral testing is performed between 9:00 AM and 6:00 PM. The experimental mice are transferred to the behavioral testing room 30 min before the first trial to allow them to habituate to the room conditions. All procedures were approved by the Committee on Ethics in the Use of Animals from Kunming Medical University (CEUA no. kmmu2021227). MA was dissolved in sterile saline to a concentration of 1 mg/ml as a stock solution. Quercetin was purchased from Sigma-Aldrich Company (Sigma-Aldrich, MO, USA) and was first dissolved in polyethylene glycol (PEG, Sigma-Aldrich), at a final concentration of 50 mg/kg in 20% PEG with 0.9% saline. Adult male C57BL/6 mice which weighed from 22 to 25 g were randomly divided into three groups (n = 12 each group): control, MA-treated, and MA + quercetin (Q)-treated. The control group received normal saline injection intraperitoneally; the MA group received escalating MA doses, as described in a previous study (Manning et al., 2016), at 5, 10, and 15 mg/kg during the first, second, and third weeks, respectively; and the MA + Q group received escalating MA doses and quercetin treatment, administered with one dose at 50mg/kg daily for 4 days during the first week and 5 days in the second week, as described in a previous study (Zhang et al., 2019). On the 22nd day, the open field (OFT) and Elevated Plus Maze (EPM) tests were used to examine the motor activity and anxiety levels, respectively. After behavioral testing, the animals were sacrificed immedietely, and subsequent experiments were conducted.

### Conditioned place preference test (CPP)

The CPP apparatus consisted of two compartments: one had black and white striped walls, a white floor, and a black ceiling; the other had black and white checkered walls, a black floor, and a white ceiling. The two compartments were separated by a removable board. Behavioral subjects were habituated for 5 min and placed in the experimental environment for adaptation for 3 days before the CPP test. Pre-test, conditioning, and a test were included in the MA CPP session. During the pre-test phase, the mice were placed in the middle of the conditioning apparatus and allowed to freely explore the full extent of the CPP apparatus for 15 min. The time spent in each chamber was measured. Mice that spent >65% (>585 s) or <35% (<315 s) of the total time (900 s) on one side were eliminated from subsequent CPP experiments (Zhou et al., 2019). Conditioning was conducted on mice confined to one chamber for 30 min, which was paired with an intraperitoneal (i.p.) MA injection on days 1, 3, 5, and 7, and on days 2, 4, and 6, the mice were confined to the other chamber for 30 min, which was paired with an i.p. saline injection. For the CPP test, mice were released from the middle part of the CPP apparatus and allowed to freely explore both chambers for 15 min. The CPP score was calculated by subtracting the time spent within the saline-paired side from that spent on the MA-paired side. Mouse behavior was analyzed using the ANY-maze video tracking system (Stoelting Co.).

### Open Field Test (OFT)

Each experimental animal was placed in the corner of the open field apparatus (50 × 50 × 40 cm^3^, SANS Co., Jiangsu, China), which consisted of a white plastic floor and wall. The OFT performance was recorded using a video camera attached to a computer and controlled by a remote device. The total distance traveled (cm) and time spent in the center area (20 × 20 cm^2^) were recorded during a 5 minute test period. After each trial, the whole open field apparatus was cleaned with 75% ethyl alcohol to efficiently remove odor to prevent any bias based on olfactory cues. The mouse behavior was analyzed using the ANY-maze video tracking system (Stoelting Co.).

### Elevated plus maze test (EPM)

The EPM consisted of two open arms (30 × 5 cm) and two enclosed arms of the same size, with 15 cm white plastic walls. The four arms were connected by a central square (5 × 5 cm) (SANS Co., Jiangsu, China). The arms were elevated 55 cm above the floor. Each experimental mouse was placed in the central square of the maze, facing one of the enclosed arms. The number of entries into each arm and the time spent in the open arms were recorded during a 5-min test period. When a mouse falls from the maze, the data are excluded. After each trial, all arms and the center area were cleaned with 75% ethyl alcohol, as previously described. All experimental data were collected and analyzed described.

### RNA preparation, library construction, and sequencing

Total RNA was isolated using RNA-Bee reagent, following the manufacturer’s protocol. RNA purity was determined using the NanoPhotometer spectrophotometer (IMPLEN, CA, USA), and the concentration was determined using the Qubit RNA Assay Kit (Life Technologies, CA, USA). Samples with RNA integrity values >7.0 were used for the following experiments, which were assessed by the RNA Nano 6000 Assay Kit of the Bioanalyzer 2100 system (Agilent Technologies, CA, USA). Sequencing libraries were prepared using the NEB Next Ultra RNA Library Prep Kit (Illumina, USA), according to the manufacturer’s recommendations, which were described in our previous research (Sun et al., 2020). In brief, mRNA was purified using poly-T oligo-attached magnetic beads, fragmentation was performed using divalent cations, and library quality was assessed on the Agilent Bioanalyzer 2100 and qPCR. The clustering of the index-coded samples was performed on an acBot Cluster Generation System using TruSeq PE Cluster Kitv3-cBot-HS (Illumina, San Diego, CA, USA). The library preparations were then sequenced on an Illumina Hiseq platform, and paired-end reads were generated.

### Quantification and differential expression analysis of mRNA

As mentioned in Sun et al. (Sun et al., 2020), the reference genome index was built, and paired-end clean reads were aligned to the reference genome using Hisat2 v2.0.5. Feature Counts v1.5.0-p3 was used to count the reads numbers. The expected number of fragments per kilobase of transcript sequence per millions (FPKM) of base pairs sequenced was calculated based on the gene length, and read counts were mapped to the gene.

Differential expression analysis (n = 3 per group) was performed using the DESeq2 R package (1.16.1). *P*-values were adjusted using the Benjamini and Hochberg approach for controlling the false discovery rate (FDR), and *p*<0.05 and log_2_ fold-change >2 were considered to be significant DEGs. DEGs enrichment in the KEGG pathways was assessed using the web tool Metascape (http://metascape.org).

### Immunofluorescence staining and imaging

Mice were deeply anesthetized and perfused with 25 ml ice-cold PBS, followed by 25 ml 4% ice-cold paraformaldehyde (PFA) in PBS. Brains were removed and dehydrated with 15% and 30% sucrose at 4°C. The fixed brains were sliced into 30-μm-thick sagittal slices using a Leica CM1950. Slices were permeabilized in 1.2% Triton X-100 in PBS for 15 min and subject to incubation in blocking solution. Slices were incubated with primary antibodies for NeuN (1:200, Abcam), glial fibrillary acidic protein (GFAP, 1:500, Abcam) for 24 h at 4°C, followed by incubation with species-matched and Alexa Fluor conjugated secondary antibodies raised in rabbit (1:5000, Invitrogen) for 2 h at room temperature. 4’,6-Diamadino-2-phenylindole (DAPI, 1:1000, Invitrogen) was incubated after secondary antibody incubation for 15 min at room temperature. Slices were mounted and coverslipped using VECTASHIELD H-1000 mounting medium and scanned on a Nikon C2 confocal microscope using NIS-Element software.

### H&E staining and Electron microscope imaging

The brain tissues were fixed in 4% paraformaldehyde (PFA) immediately after sacrifice. H&E staining was performed on 5μm paraffin sections using standard H&E staining protoccol which were described previously (Sun et al., 2020). The thin sections were made with an ultramicrotome, stained by OsO_4_. Electron microscopy and ultrastructural studies were performed on a transmission electron microscope (JEM-1400Flash).

### Measurement of ATP, mitochondrial membrane potential (MMP), and reactive oxygen species (ROS)

Intracellular ATP was determined using a firefly luciferase-based ATP assay kit (Beyotime, Beijing, China) based on a fluorescence technique. In brief, astrocytes (1 × 10^4^) were plated in 96-wells, and appropriate drug treatments were applied for 48 h. Opaque-walled 96-well plates with culture media (50 µL) were prepared. Luminescence test solution (50 µL) was added and incubated for 30 min and then measured using a luminescence microplate reader. Mitochondria-derived ATP was measured after treatment with 300 mM iodoacetic acid (IAA, Sigma-Aldrich, MO, USA). IAA was added to half of the wells and the cells were then incubated at 37 °C and 5% CO_2_ for 60 min. After 30 min, 1 mM oligomycin was added to half of the wells containing IAA and incubated for 30 min to abolish all ATP production and confirm that the ATP levels in the presence of IAA were produced by the mitochondrial ATP synthase. Subsequently, the media was removed from the wells and cellular ATP was measured using ATP assay kit (mentioned before).

JC-1 assay was conducted to analyze the MMP using the JC-1 mitochondrial membrane potential assay kit (Beyotime, Beijing, China). In brief, astrocytes (2 × 10^5^) were plated in 6-well plates and, after appropriate drug treatments, stained with 10 µM JC-1 for 20 min. JC-1 exhibits double fluorescence staining, either as red fluorescent J-aggregates (530 nm excitation/590 nm emission, as P3) at high potentials or as green fluorescent J-monomers (490 nm excitation/530 nm emission, as P2) at low potentials; Flow cytometric analysis was performed by fluorescence-assisted cell sorting (FACS) after JC-1 staining detected changes in MMP and the value was calculated. The relative proportion of red and green fluorescence was used as an index of change in membrane potential.

For ROS generation measurements, primary astrocytes were plated in 96-wells, and appropriate drug treatments were applied for 48 h. Then, cells were loaded with the ROS probe 70-dichlorodihydrofluorescein diacetate (DCFDA, 2 µM) for 40 min at RT in the dark. The mROS assay was conducted using the MitoSOX red mitochondrial superoxide indicator (Mercury Drive, Sunnyvale, CA). Cells were measured by FACS at 488 nM excitation.

### Oxygen consumption rate (OCR) analysis

Astrocytes were plated at 7.5 × 10^4^/well in seahorse assay plates and treated with their respective treatments at 37°C and 5% CO_2_ and for 24 h. Mitochondrial oxygen consumption rate (OCR) was measured by extracellular flux (XF) assay (Seahorse XFp analyzer, Agilent Technologies, Santa Clara, CA), according to manufacturer’s procedures. Briefly, cells were incubated in a CO_2_-free environment for 1 h, and OCR was measured every 3 min for the next 90 min. First, OCR was acquired in basal conditions (20 mM glucose), followed by in the presence of 1.5 µM oligomycin (ATP synthase inhibitor), with 3.5 µM carbonyl cyanide-p-trifluoromethoxy phenylhydrazone (FCCP), and, finally, with 0.5 µM rotenone/antimycin A.

### Quantification of gene expression assay

Quantification of gene expression was performed by ABI 7500 Sequence Detection System (Applied Biosystems, Foster City, CA, USA). The qPCR assays were performed as described in our previous study (Chen et al., 2016). The primers used for qPCR are shown in **Table S1**. The mRNA levels were determined by qPCR in triplicate for each of the independently prepared RNA samples, and mRNA levels were normalized against the levels of glyceraldehyde 3-phosphate dehydrogenase (*GAPDH)* expression.

### Western blotting analysis

Total protein was extracted from astrocytes or brain tissue and quantified by bicinchoninic acid (BCA) protein assay kit (Pierce, USA) and used for immunoblotting analysis, as described previously. Briefly, the blot was incubated with a specific primary antibody overnight (anti-GFAP, 1:1000, Abcam; anti-MFN2, 1:200, Abcam; anti-OPA1, 1:200, Abcam; anti-DRP1, 1:200, Abcam), followed by incubation with HRP-conjugated secondary antibody. The bands were detected with a chemiluminescence detection kit (Millipore Co., MA, USA) and scanned using the iBright FL1500 chemiluminescence imaging system (Thermo Fisher Scientific, USA).

### Statistical analysis

The results are presented as the mean ± standard error of the mean (SEM). For comparisons between two or multiple groups, the Student’s t-test or one-way analysis of variance (ANOVA) analysis was conducted, respectively. Significance is indicated by asterisks: \**p*<0.05, \*\**p*<0.01, \*\*\**p*<0.001.

## RESULTS

### Repeated MA administrations induce anxiety-like behavior in mice

To identify an optimal concentration for the generation of an MA-addicted mouse model, we tested three different MA concentrations (2.5, 5, and 10 mg/kg, i.p.). Mice are trained to associate the MA reward with the paired context during training for the MA CPP (**Figure 1A**). A preference for the MA-paired side indicates the expression of a reward–context associated memory, which was assessed by measuring the time that an animal spent on the MA-paired side in the CPP apparatus. After four sessions of MA CPP training, mice treated with 5 mg/kg MAshowed a significant preference for the MA-paired side. The 2.5 mg/kg MA group (**Figure 1B**, *p* < 0.05) also showed a significant preference for the MA-paired side. However, 6 of 8 mice died in the 10 mg/kg MA-treated group. Therefore, we chose 5 mg/kg MA as the initial concentration to establish a subsequent mouse model (**Figure 1C**).

**Figure 1.**
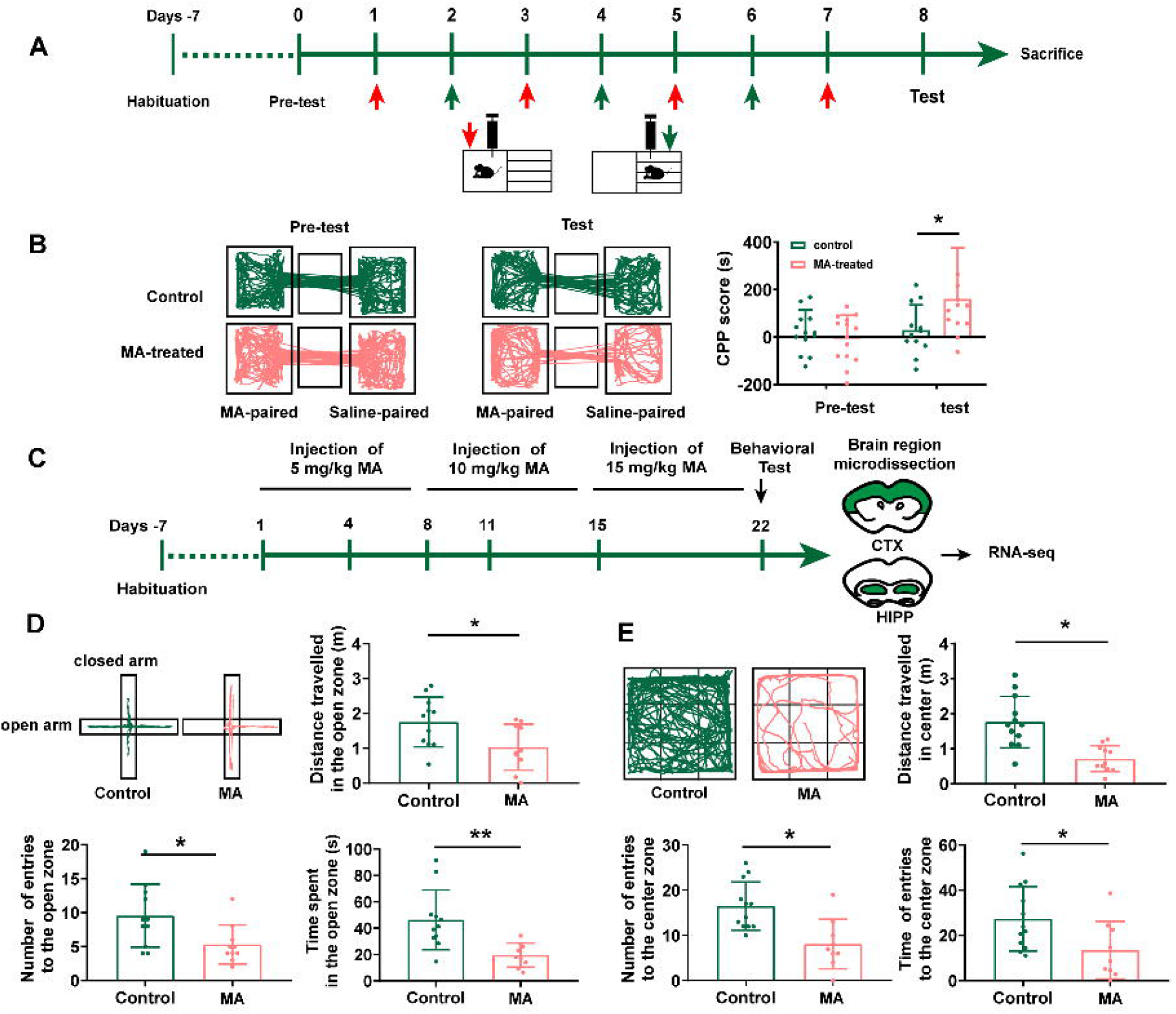
Repeated MA administrations induce anxiety-like behavior in mice. A. Timeline of the Conditional place preference (CPP) experimental procedure. B. The movement trajectory of the two groups of mice across compartments (left); CPP scores were assessed as the difference of time spent in the drug-paired compartment between the post and pre-conditioning phases (right). C. Timeline of the model establishment and sample collection. D and E. Anxiety assessment with the elevated plus maze (EPM) and open field test (OFT). Student- t-tests, \**p* < 0.05, \*\**p* < 0.05.

Mice with 3-week MA-treated mice demonstrated a fear of entering the open arms of the EPM test (**Figure 1D**) with fewer numbers of entries to open zone and less time(\*\**p* < 0.01) and distance(\**p* < 0.05) spent in the open zone. Consistently, MA treatments also displayed decreased locomotor activity and spent significantly less time in the center of the OFT than control mice (**Figure 1E**, \**p* < 0.01), suggesting that we successfully generated an MA-treated mouse model with anxiety-like behaviors.

### RNA-seq revealed differentially expressed genes in the HIPP in the MA-treated mouse model

To better elucidate the molecular features of MA-induced anxiety-like behaviors in mice, we performed RNA-seq to investigate DEGs between control and chronic MA-treated mice in HIPP. Hierarchical clustering analysis was performed and displayed in **Figure 2A**. Forteen upregulated genes were identified between the control and MA-treated groups, which were displayed in **Supplementary Table 1**.

**Figure 2.**
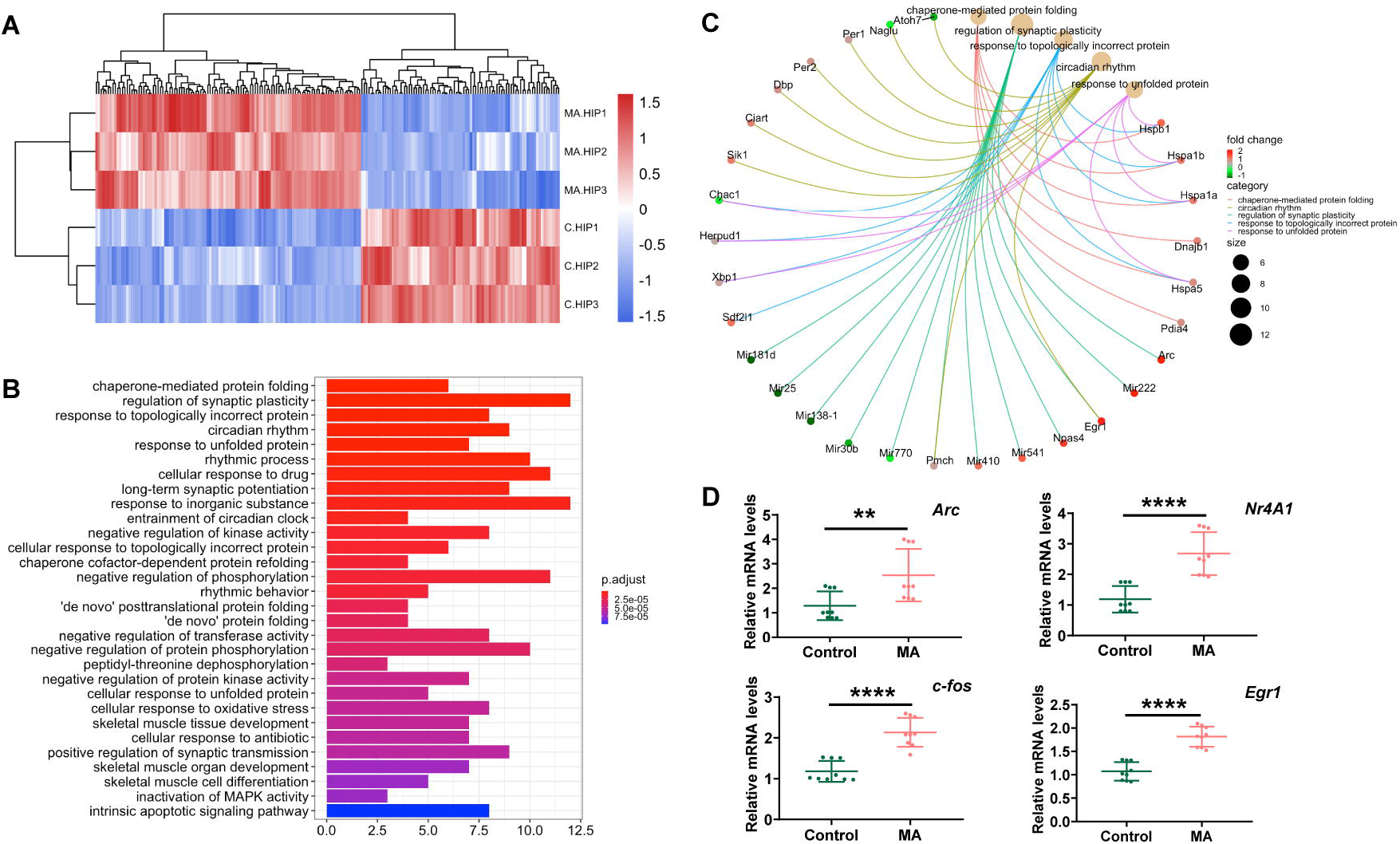
RNA-seq revealed differentially expressed genes in the HIPP in the MA-treated mouse model. A. Hierarchical clustering analysis of RNAs with altered expression between the two groups (p < 0.05, fold change > 2). Red strip, high relative expression; blue strip, low relative expression. Color intensity reflects the degree of expression increase or decrease. B. Thirty most enriched KEGG classifications of assembled differential genes in HIPP. C. The genes implicated by the Top 5 KEGG signalling pathways in HIPP. D. qPCR validation for RNA-seq data.

To investigate the altered signaling pathways involved in mice presenting with MA-induced anxiety-like behavior, we performed KEGG enrichment analyses. DEGs between the control and MA-treated groups were mainly enriched in signaling pathways associated with protein folding (chaperone-mediated protein folding, response to unfolded protein, chaperone cofactor-dependent protein refolding, ‘de novo’ posttranslational protein folding, ‘de novo’ protein folding, cellular response to unfolded protein), synaptic plasticity and synaptic transmission (regulation of synaptic plasticity, positive regulation of synaptic transmission), rhythmic processes (regulation of synaptic plasticity, positive regulation of synaptic transmission), protein phosphorylation (negative regulation of phosphorylation, negative regulation of protein phosphorylation, peptidyl-threonine dephosphorylation), oxidative stress, and the intrinsic apoptotic signaling pathway (**Figure 2B**). As Mitochondria couple with the endoplasmic reticulum (ER) are important in protein folding (Marchi, et al. 2014). These results suggested mitochondrial and ER play an important role in MA-induced anxiety and might be promising therapeutic targets.

Furthermore, as immediate early genes (IEGs) have been reported to be involved in methamphetamine use and the development of neuropsychiatric disorders (McCoy et al., 2011; Gallitano et al., 2020), we used three samples from each group to validate four selected DEGs classified as IEGs (*Fos, Egr1, Arc* and *Nr4a1*). The results revealed that, in general, MA upregulated all the four IEGs identified, in accordance with the RNA-seq results (**Figure 2D**).

### Administration of quercetin ameliorates anxiety-like behaviors in an MA mouse model

Quercetin has been reported to mitigate anxiety-like symptoms in a lipopolysaccharide-induced mouse model of anxiety (Lee et al., 2020; Samad et al., 2018) and has been reported as a mediator of mitochondrial function and ER stress (Khan et al., 2016). Therefore, we evaluated the effects of quercetin on MA-induced behavioral phenotypes and the role played by quercetin in mitochondrial functional modifications in the present study (**Figure 3A**).

**Figure 3.**
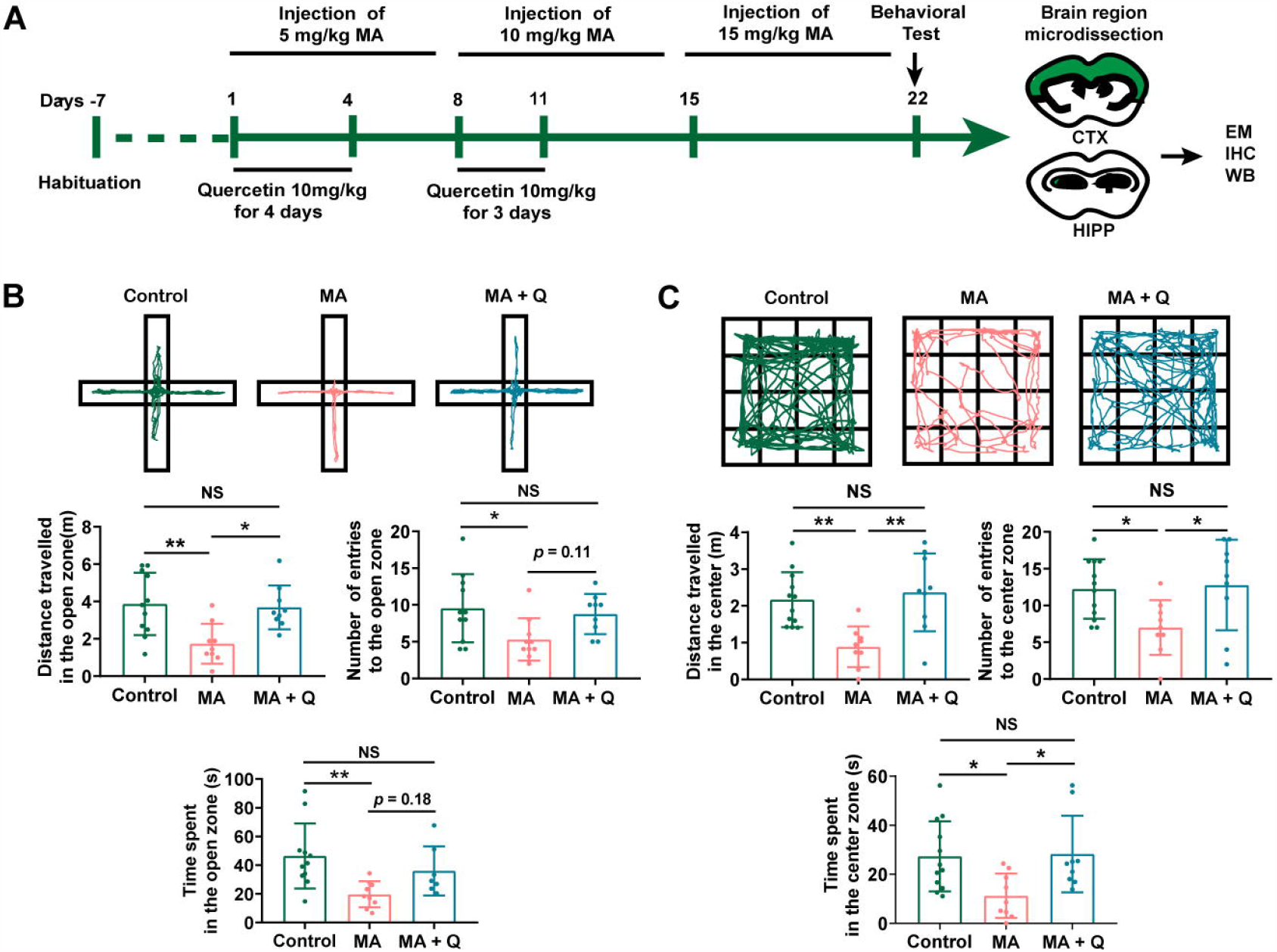
Administration of quercetin ameliorates anxiety-like behaviors in an MA mouse model. A. Timeline of the model establishment and sample collection. B and C. Anxiety assessment with the elevated plus maze (EPM) and open field test (OFT). Student t-tests, \**p* < 0.05, \*\**p* < 0.05.

In the EPM test, the time spent and the number of entries into the open arms were significantly reduced in the MA-treated group compared with those in the control group (**Figure 3B**, *p* < 0.05). Conversely, when compared with the control group, the time spent and number of entries into the closed arms were significantly elevated in the MA-treated group (**Data not shown**). However, mice treated with MA□and quercetin showed the significant restoration of the time spent and the number of entries into the open arms compared with those for the single MA-treated group (**Figure 3B**, *p* < 0.05), with no significant differences observed between the control group and the MA□+□Q group (**Figure 3B**, *p* > 0.05).

The OFT results revealed that a single administration of MA significantly reduced the number of times MA-treated mice crossed in the central zone compared to the saline-treated group (**Figure 3C**, *p* < 0.05), but no significant difference in the number of crossings in the peripheral zone were observed (**Data not shown**). Quercetin combined with MA treatment significantly enhanced the number of central zone crossings compared with the single MA-treated group (**Figure 3C**, *p* < 0.05).

### Quercetin alleviates neuronal injury and decreases astrocyte activation induced by MA *in vivo*

Most of previous researches have focused on MA-induced neuronal injury. Nevertheless astrocytes have been shown to participate in anxiety development through the synaptic pruning of neurons (Çalışkan et al., 2020), and they were reported to be activated in an MA-treated animal model, resulting in morphological and phenotypic changes (Zhou et al., 2019). To assess the change of neurons and astrocytes in anxiety induced by MA, we first assessed the numbers of neuron by immunostaining for NeuN and pathological alterations by H&E staining. And then to evaluate astrocytes activation by immunostaining and western blotting for GFAP. The results demonstrated MA administration markedly decreased the numbers of neurons (**Figure 4A**), numerous impaired neurons with karyopyknosis, cell gaps, and debris in dentate gyrus (**Figure 4B**). Furthermore, after MA treatment GFAP expression increased (**Figure 4G**, *p* < 0.05) with a larger area of astrocytes (**Figure 4C-E**, *p* < 0.01). Quercetin rescued the decreasing of neuron numbers and neuron damage, and attenuated the activation of astrocytes (**Figure 4C-E**, *p* < 0.01), suggesting that MA treatment facilitated neuronal loss and astrocytes activation, whereas quercetin reversed these cellular phenotypes.

**Figure 4.**
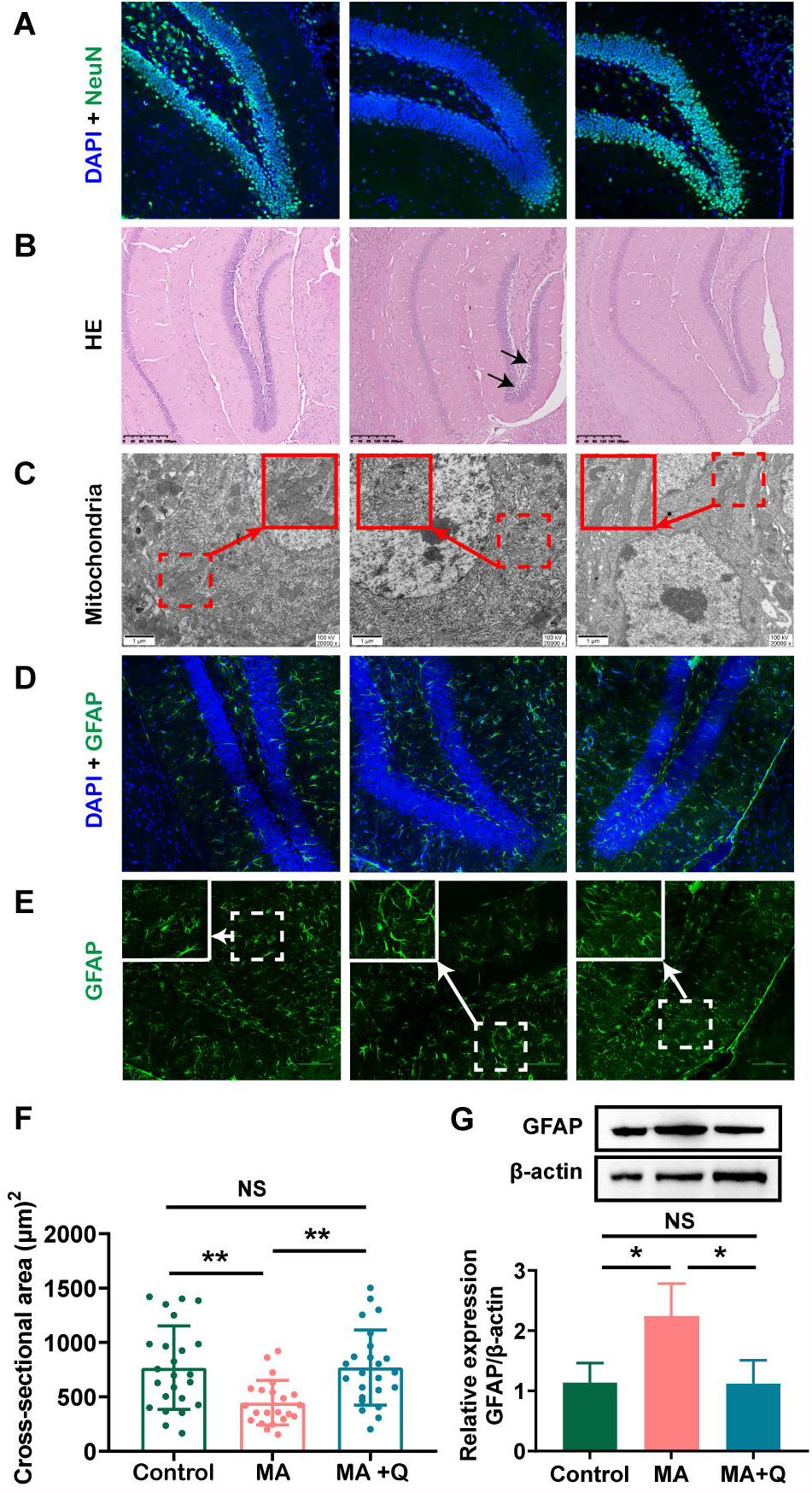
Quercetin alleviates neuronal injury and decreases astrocyte activation in HIPP induced by MA *in vivo*. A. Immunofluorescence was performed with DAPI (blue) and anti-NeuN (green). B. Hippocampus sections with H&E staining. C. Electron microscopy analysis (magnification, ×20,000). Areas in red boxes are magnified (Insets). D. Immunofluorescence was performed with DAPI (blue) and anti-GFAP (green). E-G. GFAP intensity and area were quantified to evaluate astrocytes reactivity. E. Representative image of GFAP labels astrocytes. F. Area of astrocytes (µm2/cell) in each group, \**p* < 0.05, \*\**p* < 0.01. G. Representative band pattern of the WB of different treatment of astrocytes using antibodies for GFAP and β-Actin (left); summary bar graphs of GFAP and β-Actin levels in different group (right).

We also evaluated mitochondrial morphology in the HIPP using electron microscopy. The control group presented with relatively long, tubular mitochondria, whereas the mitochondria in the HIPP of MA-treated mice were relatively more fragmented and swollen, with the loss of cristae (**Figure 4D**), indicating MA-induced damage to mitochondria in HIPP. Quercetin treatment was able to reinstate mitochondrial morphology toward long tubular mitochondria with intact cristaein MA-treated mice(**Figure 4D**).

### Quercetin ameliorates mitochondrial dysfunction and aberrant morphology in astrocytes *in vitro*

To assess the effect of MA on mitochondria *in vitro*, we conducted experiments in astrocytes. We found that when astrocytes were exposed to MA, tubular mitochondria shortened in length and swelled in width, forming large, spherical structures (**Figure 5A**), and similar morphological changes in mitochondria were viewed using mitotracker (**Figure 5B**). quercetin treatment rescued abnormal mitochondrial morphology in astrocytes treated with MA (**Figure 5A-B**).

**Figure 5.**
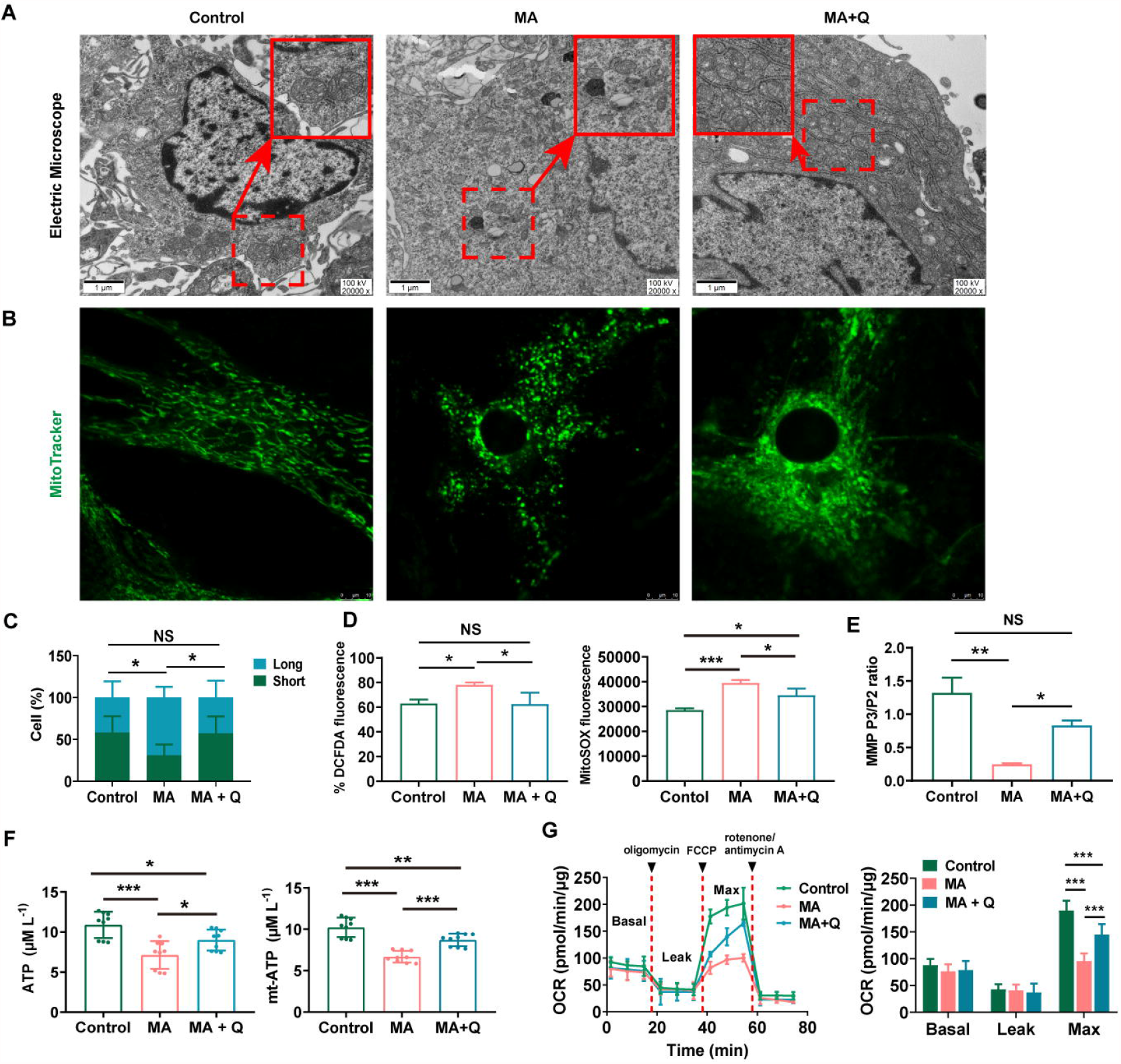
Quercetin ameliorates mitochondrial dysfunction and aberrant morphology in astrocytes *in vitro*. A. Electron microscopy analysis (magnification, ×20,000). Areas in red boxes are magnified (Insets). B. Representative image of astrocytic mitochondria. Alive astrocytes were incubated with MitoTraker Green as a probe for mitochondria in each group. The percentage of astrocytes for short-shape (blue) and long-shape (green) mitochondria in different group is presented as histograms in C. D. Total ROS (left) and mitochondria-derived ROS (right) production quantification by flow cytometry in each group. D. Quantification of the mitochondrial membrane potential (MMP). F. Quantification of the total (left) and mitochondria-derived ATP (right) by using an ATP quantification kits. G. An analysis of O2 consumption in astrocytes. The Agilent SeahorseXFe24 analyzer measures OCR at basal and after injection of oligomycin (3.5□μM), FCCP (4□μM), and antimycin A (1□μM)/rotenone (1□μM) for three measurement cycles at each step (left). Basal, ATP-linked, maximal, and reserve capacity OCR in each group. \**p* < 0.05, \*\**p*□<0.01, \*\*\**p*□<0.001 vs. control, as determined by Student’s t test.

In addition, mitochondria have been considered to serve as the intracellular source of ROS in animal cells, and when confronting with oxidative stress, mitochondrion is the primary target attacked by ROS, and also produces excessive amounts of ROS due to the damage to enzymes in the electron transport chain. Therefore, we first evaluated the total ROS by DCFDA and the mitochondrial ROS by MitoSOX staining and quantified the fluorescence intensity in MA-treated astrocytes with or without quercetin treatment. The results showed that both the total ROS and mitochondrial ROS were markedly increased in MA treated group, which were rescued by quercetin (**Figure 5D**).

A series of redox reactions creates an electrochemical gradient through the mitochondrial electron transport chain, which drives the synthesis of ATP (Mitchell et al., 1961) and generates the MMP. In this study, we evaluated the ability to generate total ATP and mitochondria-derived, and measured the MMP to represent mitochondrial function. The results revealed that the quercetin supplementation of MA-treated astrocytes markedly elevated both the total and mitochondria-derived ATP levels (**Figure 5E**) and reversed the observed decrease in MMP in MA-treated astrocytes (**Figure 5F**). We used an XF24 metabolic bioanalyzer to assess the effects of quercetin supplementation on OCR and glycolytic flux rates in MA-treated astrocytes and found that quercetin markedly reinstated the decrease in the OCR induced by MA (**Figure 5G**).

## DISCUSSION

In the present study, we found that quercetin attenuated anxious symptoms and pathology, and modified mitochondria in both a mouse model and cultured astrocytes treated with MA. These findings, for the first time, suggested that quercetin could inhibit the progression of anxiety induced by MA by modulating mitochondrial function and morphology in the central nervous system. These results indicate that quercetin represents a potential therapeutic medicine for future development and support the hypothesis that changes in mitochondrial function mediate anxiety progression.

We found that MA-induced anxiety-like behavior in mice, as previously described (Ru et al., 2019). Consistent with our findings, Iwazaki et al (Iwazaki et al., 2008)have demonstrated synaptic plasticity- and synaptic transmission-, oxidative stress-, and intrinsic apoptotic-related signaling pathway were dysregulated, implying that mitochondria as a target organelle in MA-induced anxiety. Mitochondria, which are capable of generating secondary ROS and amplifying initial oxidative insults, are both a source and a target of oxidative stress (Song et al., 2017). Mitochondrial dysfunction can also be triggered by the accumulation of misfolded proteins (Saito et al., 2018) and failed to maintain synaptic ion homeostasis and synaptic plasticity due to decreased ATP production and overloaded Ca^2+^ concentrations (Fang et al., 2015). Furthermore, mitochondria function which play an important role in apoptosis via the intrinsic apoptotic program was identified to be disregulated in frontal cortex in MA-treated rat (Iwazaki et al., 2008). The parameters about mitochondria function were abberrant in our results.Based on these, we speculate that mitochondria represent a primary target organelle of MA treatment in the HIPP.

Furthermore, four identified IEGs (*Fos, Egr1, Arc*, and *Nr4a1)* in our animal model, which were reported to change in chronic and acute MA-treated mice (Cheng et al., 2015; Akiyama et al., 2008; McCoy et al., 2011), were validated by qPCR in the transcriptomics analysis of chronic MA-treated mouse brains. The changes observed in these genes by qPCR and western blot were consistent with our RNA-seq results. Nr4a1 (also known as TR3 or NGFI-B) is an orphan member of the nuclear receptor superfamily, which migrates from the nucleus to the mitochondria, where it binds to Bcl-2 to induce apoptosis and cause the release of cytochrome c (Liu et al., 2008). According to previous reports, Nr4a1 can evoke cellular oxidative stress and disrupt ATP generation (Zhang et al., 2018), and the expression and activity of Nr4a1 are sustained by chronic stress in animal models and in human studies of neuropathologies sensitive to the buildup of chronic stress (Jeanneteau et al., 2018). Nr4a1 has been deemed to be important for regulating metabolic and morphological aspects of neuronal functions by modifying the expression of several mitochondrial regulatory genes, including *Mfn1, Mfn2, Fis1* and *OPA1* (Jeanneteau et al., 2018), which can result in mitochondrial dysfunction and behavioral phenotypes. Thus, we speculated that quercetin might modulate genes associated with mitochondrial function via downregulating Nr4a1. We then detected Mfn2, OPA1 and Drp1 expression at both the protein and mRNA levels after quercetin treatment. As anticipated, quercetin reversed the MA-induced changes in Nr4a1, OPA1 and Drp1 expression at both the protein and mRNA levels (**Supplementary Figure 1**), but failed to rescued Mfn2 expression downregulated by MA-treated. Overall, quercetin was capable of targeting mitochondria to alleviate neuronal impairments and the anxiety-like behaviors, probably through the Nr4a1–OPA1/ Drp1-mitochondrial morphology and functional pathway.

In this article, we demonstrated that chronic MA exposure resulted in the dysregulation of mitochondrial morphology and mitochondrial dysfunction, suggesting that the modification of mitochondrial function and morphology might represent a potential strategy for alleviating anxiety-like behaviors induced by MA. Accumulating data has highlighted the contributions of brain mitochondria and bioenergetics to the development of psychiatric disorders and stress-related pathologies (Manji et al., 2012; Einat et al., 2005). The fundamental role played by mitochondria in the synthesis of the primary excitatory neurotransmitter (glutamate) and the inhibitory neurotransmitter (γ-aminobutyric acid, GABA) suggest that the mitochondrial adaptations observed in the context of anxiety may contribute to an imbalance in neural excitation and inhibition, which is thought to underlie several neuropsychiatric disorders (Filiou et al., 2019). Mitochondria-targeting drugs have been reported to show good outcomes in anxiety-related studies in both humans and rodents (Filiou et al., 2019). Quercetin is a bioactive compound with diverse pharmacologic effects that have been reported to exert several beneficial effects, including neuroprotective effects (Costa et al., 2016), the regulation of the sleep–wake cycle (Kambe et al., 2010), and the optimization of mitochondrial function (Houghton et al., 2018). In this study, quercetin supplementation significantly mitigated MA-induced anxiety-like behavior by improving mitochondrial morphology and function, alleviating neuronal injury, both *in vivo* and *in vitro*. In summary, mitochondrial damage and intracellular ROS production are closely related (Cao et al., 2017) and metabolic disturbance (Li et al., 2015), eventually leading to mitochondrial dysfunction (Xie et al., 2016). The damaged mitochondria accumulation induced, synaptic loss, and neuron apoptosis (de la Mata et al., 2017), underscoring the importance of mitochondria in the development of psychotic disorders. Here, we also identified the novel pharmacological efficacy of quercetin in the treatment of anxiety induced by MA.

MA exposure facilitated mitochondrial dysfunction and morphological abnormalities (Valian et al., 2019), which can trigger various innate immune signaling pathways in a cell-intrinsic or -extrinsic manner (Bader et al., 2020), resulting in the release of inflammatory factors. In this study, we found that MA treatment accelerated astrocyte activation, accompanied by the elevation of interleukin (IL)-1β, IL-6, and tumor necrosis factor (TNF)α in the HIPP and MA-treated astrocytes (**Supplementary Figure 2**). Astrocytes serve as crucial regulators of the innate and adaptive immune responses, which play critical roles in neuroinflammation (Colombo et al., 2016). According to the previous research, astrocytes can dispose of and recycle damaged mitochondria released by neurons, and release healthy extracellular mitochondrial particles to neuron (Hayakawa et al., 2016). As a major source of glycogen and lactate, astrocytes provided neurons with additional energy and play key metabolic roles in the CNS (Shirakawa et al., 2010). Moreover, astrocyte dysfunction has been shown to facilitate the pathogenesis of neurological and psychiatric disorders (Tian et al., 2018). Quercetin is a bioactive compound with antioxidant and anti-inflammatory properties (Khan et al., 2016) and was able to rescue mitochondrial dysfunction and morphological abnormalities and reduce the concentration of IL-1β, IL-6, and TNFα in astrocytes treated with MA (**Figure 5 and Supplementary Figure 2**). Malfunctioning mitochondria in astrocytes trigger various innate immune signaling pathways in a cell-intrinsic or -extrinsic manner (Bader et al., 2020), resulting in the release of inflammatory factors. These results suggested that MA treatment accelerated astrocyte activation and facilitated the release of inflammatory factors, which can trigger neuronal apoptosis and synaptic loss (Garwood et al., 2011; Çalışkan et al., 2020). Moreover, the excessive accumulation of damaged mitochondria, combined with astrocyte activation and the increase in inflammatory factors, induces neurotoxicity and promotes neuronal apoptosis (Nicholls et al., 2004), which initiates a vicious cycle that aggravates the anxiety process (Çalışkan G et al., 2020; Garwood et al., 2011). These findings also confirmed the neuroprotective activity of quercetin, through rebalancing neuroinflammation levels and modulating astrocytes activation.

## CONCLUSION

In conclusion, our study indicated that MA damaged brain cells, including neurons and astrocytes, in the HIPP by altering mitochondrial metabolism, energy, and morphology in astrocytes, which can cause the development of neuropsychiatric disorders, such as anxiety. The oral supplementation of quercetin neutralized the neuropsychiatric status of MA-treated mice, and our findings support the potential of quercetin as a candidate agent for alleviating MA-induced anxiety and support the hypothesis that mitochondria mediate anxiety progression. Further study is necessary to illustrate the contributions and mechanisms of mitochondria in the progression of anxiety induced by MA.

## DATA AVAILABILITY STATEMENT

The original contributions presented in the study are included in the article and Supplementary Material, further inquiries can be directed to the corresponding authors.

## ETHICS STATEMENT

All our studies using Kunming mice were approved by the the Committee on Ethics in the Use of Animals from Kunming Medical University (CEUA no. kmmu2021227) and were performed in accordance with Yunnan Administration Rule of Laboratory Animal.

## AUTHORS CONTRIBUTIONS

FC and JY designed the experiments, participated in the data analysis, and wrote the manuscript; JS and LZ performed the sequence alignment and bioinformatics, YZ, YD, and WT performed the animal experiments, FC, CC, and ZZ performed the cell experiments. HW, YX, and MZ participated in data analysis. JY and KW conceived of the study, supervised the project, and participated in the drafting of the manuscript. All authors read and approved the final manuscript.

## ACKNOWLEDGEMENTS

We would like to thank Professor Bai Jie and Kong Xiangyang for the technical support of behavioral test. Also, we thank the technical support of Dr Li Shaoyou for the Flow Cytometry test.

## REFERENCES

Akiyama K, Isao T, Ide S, Ishikawa M, Saito A. mRNA expression of the Nurr1 and NGFI-B nuclear receptor families following acute and chronic administration of methamphetamine. Prog Neuropsychopharmacol Biol Psychiatry. 2008;32(8):1957–66.

Andres S, Pevny S, Ziegenhagen R, Bakhiya N, Schäfer B, Hirsch-Ernst KI, Lampen A. Safety Aspects of the Use of Quercetin as a Dietary Supplement. Mol Nutr Food Res. 2018;62(1).

Bader V, Winklhofer KF. Mitochondria at the interface between neurodegeneration and neuroinflammation. Semin Cell Dev Biol. 2020;99:163–171.

Çalışkan G, Müller A, Albrecht A. Long-Term Impact of Early-Life Stress on Hippocampal Plasticity: Spotlight on Astrocytes. Int J Mol Sci.2020;15;21(14):4999.

Cao JJ, Tan CP, Chen MH, Wu N, Yao DY, Liu XG, Ji LN, Mao ZW. Targeting cancer cell metabolism with mitochondria-immobilized phosphorescent cyclometalated iridium (iii) complexes. Chem Sci. 2017;8(1):631–640.

Chang L, Alicata D, Ernst, T, Volkow N. Structural and metabolic brain changes in the striatum associated with methamphetamine abuse. Addiction, 2017;102(Suppl 1):16–32.

Chen FR, Zhu LP, Cai L, Zhang JW, Zeng XQ, Li JS, Su Y, Hu QH. A stromal interaction molecule 1 variant up-regulates matrix metalloproteinase-2 expression by strengthening nucleoplasmic Ca(2+) signaling. BBA-Molecular Cell Research, 2016;1863(4):617–29.

Cheng MC, Hsu SH, Chen CH. Chronic methamphetamine treatment reduces the expression of synaptic plasticity genes and changes their DNA methylation status in the mouse brain. Brain Res. 2015;1629:126–34.

Chiang M, Lombardi D, Du J, Makrum U, Sitthichai R, Harrington A, Shukair N, Zhao M, Fan X. Methamphetamine-associated psychosis: Clinical presentation, biological basis, and treatment options. Hum Psychopharmacol. 2019;34(5):e2710.

Colombo E, Farina C. Astrocytes: Key Regulators of Neuroinflammation. Trends Immunol. 2016;37(9):608–620.

Costa LG, Garrick JM, Roquè PJ, Pellacani C. Mechanisms of Neuroprotection by Quercetin: Counteracting Oxidative Stress and More. Oxid Med Cell Longev. 2016;2016:2986796.

de la Mata M, Cotán D, Oropesa-Ávila M, Villanueva-Paz M, de Lavera I, Álvarez-Córdoba M, Luzón-Hidalgo R, Suárez-Rivero JM, Tiscornia G, Sánchez-Alcázar JA. Coenzyme Q10 partially restores pathological alterations in a macrophage model of Gaucher disease. Orphanet J Rare Dis. 2017;12(1):23.

Dhiman P, Malik N, Sobarzo-Sánchez E, Uriarte E, Khatkar A. Quercetin and Related Chromenone Derivatives as Monoamine Oxidase Inhibitors: Targeting Neurological and Mental Disorders. Molecules. 2019;24(3):418.

Einat H, Yuan P, Manji HK. Increased anxiety-like behaviors and mitochondrial dysfunction in mice with targeted mutation of the Bcl-2 gene: Further support for the involvement of mitochondrial function in anxiety disorders. Behavioural Brain Research. 2005;165:172–180.

Fang D, Wang Y, Zhang Z, Du H, Yan S, Sun Q, Zhong C, Wu L, Vangavaragu JR, Yan S, Hu G, Guo L, Rabinowitz M, Glaser E, Arancio O, Sosunov AA, McKhann GM, Chen JX, Yan SS. Increased neuronal PreP activity reduces Aβ accumulation, attenuates neuroinflammation and improves mitochondrial and synaptic function in Alzheimer disease’s mouse model. Human molecular genetics. 2015;24(18):5198–5210.

Filiou MD, Sandi C. Anxiety and Brain Mitochondria: A Bidirectional Crosstalk. Trends Neurosci. 2019;42(9):573–588.

Gallitano AL. Editorial: The Role of Immediate Early Genes in Neuropsychiatric Illness. Front Behav Neurosci. 2020;21;14:16.

Garwood CJ, Pooler AM, Atherton J, Hanger DP, Noble W. Astrocytes are important mediators of Aβ-induced neurotoxicity and tau phosphorylation in primary culture. Cell Death Dis. 2011;2(6):e167.

Glasner-Edwards S, Mooney LJ, Marinelli-Casey P, Hillhouse M, Ang A, Rawson R, Methamphetamine Treatment Project Corporate A. Anxiety disorders among methamphetamine dependent adults: association with post-treatment functioning. American Journal on Addictions. 2010;19(5):385–390.

Glasner-Edwards S, Mooney LJ. Methamphetamine psychosis: epidemiology and management. CNS Drugs. 2014;28(12):1115–1126.

Golsorkhdan SA, Boroujeni ME, Aliaghaei A, Abdollahifar MA, Ramezanpour A, Nejatbakhsh R, Anarkooli IJ, Barfi E, Fridoni MJ. Methamphetamine administration impairs behavior, memory and underlying signaling pathways in the hippocampus. Behav Brain Res. 2020;379:112300.

Hayakawa K, Esposito E, Wang X, Terasaki Y, Liu Y, Xing C, Ji X, Lo EH. Transfer of mitochondria from astrocytes to neurons after stroke. Nature. 2016;535(7613):551–5.

Hellem TL. A Review of Methamphetamine Dependence and Withdrawal Treatment: A Focus on Anxiety Outcomes. J Subst Abuse Treat. 2016; 71:16–22.

Homer BD, Solomon TM, Moeller RW, Mascia A, DeRaleau L, Halkitis PN. Methamphetamine abuse and impairment of social functioning: a review of the underlying neurophysiological causes and behavioral implications. Psychol Bull. 2008;134:301–310.

Houghton MJ, Kerimi A, Tumova S, Boyle JP, Williamson G. Quercetin preserves redox status and stimulates mitochondrial function in metabolically-stressed HepG2 cells. Free Radic Biol Med. 2018;129:296–309.

Huang RR, Zhang Y, Han B, Bai Y, Zhou RB, Gan GM, Chao J, Hu G, Yao HH. Circular RNA HIPK2 regulates astrocyte activation via cooperation of autophagy and ER stress by targeting MIR124-2HG. Autophagy. 2017;13(10):1722–1741.

Iwazaki T, McGregor IS, Matsumoto I. Protein expression profile in the amygdala of rats with methamphetamine-induced behavioral sensitization. Neurosci Lett. 2008;18;435(2):113–9.

Jeanneteau F, Barrère C, Vos M, De Vries CJM, Rouillard C, Levesque D, Dromard Y, Moisan MP, Duric V, Franklin TC, Duman RS, Lewis DA, Ginsberg SD, Arango-Lievano M. The Stress-Induced Transcription Factor NR4A1 Adjusts Mitochondrial Function and Synapse Number in Prefrontal Cortex. J Neurosci. 2018;7;38(6):1335–1350.

Sun JX, Chen FR, Chen C, Zhang ZR, Zhang ZY, Tian WW, Yu JH, Wang KH. Intestinal mRNA expression profile and bioinformatics analysis in a methamphetamine-induced mouse model of inflammatory bowel disease. Ann Transl Med. 2020;8(24):1669.

Kambe D, Kotani M, Yoshimoto M, Kaku S, Chaki S, Honda K. Effects of quercetin on the sleep-wake cycle in rats: involvement of gamma-aminobutyric acid receptor type A in regulation of rapid eye movement sleep. Brain Res. 2010;1330:83–8.

Khan I, Paul S, Jakhar R, Bhardwaj M, Han J, Kang SC. Novel quercetin derivative TEF induces ER stress and mitochondria-mediated apoptosis in human colon cancer HCT-116 cells. Biomed Pharmacother. 2016;84:789–799.

Kohno M, Loftis JM, Huckans M, Dennis LE, McCready H, Hoffman WF. The relationship between interleukin-6 and functional connectivity in methamphetamine users. Neuroscience Letters. 2018;677:49–54.

Kosari-Nasab M, Shokouhi G, Ghorbanihaghjo A, Mesgari-Abbasi M, Salari AA. Quercetin mitigates anxiety-like behavior and normalizes hypothalamus-pituitary-adrenal axis function in a mouse model of mild traumatic brain injury. Behav Pharmacol. 2019;30(2 and 3-Spec Issue):282–289.

Lee B, Yeom M, Shim I, Lee H, Hahm DH. Protective Effects of Quercetin on Anxiety-Like Symptoms and Neuroinflammation Induced by Lipopolysaccharide in Rats. Evid Based Complement Alternat Med. 2020;2020:4892415.

Li P, Wang B, Sun F, Li Y, Li Q, Lang H, Zhao Z, Gao P, Zhao Y, Shang Q, Liu D, Zhu Z. Mitochondrial respiratory dysfunctions of blood mononuclear cells link with cardiac disturbance in patients with early-stage heart failure. Sci Rep. 2015;5:10229.

Liu J, Zhou W, Li SS, Sun Z, Lin B, Lang YY, He JY, Cao X, Yan T, Wang L, Lu J, Han YH, Cao Y, Zhang XK, Zeng JZ. Cancer Res. 2008;68(21):8871–80.

Manji H, Kato T, Di Prospero NA, Ness S, Beal MF, Krams M, Chen G. Impaired mitochondrial function in psychiatric disorders. Nat Rev Neurosci. 2012;13:293–307.

Manning EE, Halberstadt AL, van den Buuse M. BDNF-Deficient Mice Show Reduced Psychosis-Related Behaviors Following Chronic Methamphetamine. Int J Neuropsychopharmacol. 2016;19(4):pyv116.

Marchi S, Patergnani S, Pinton P. The endoplasmic reticulum-mitochondria connection: one touch, multiple functions. Biochim. Biophys. Acta. 2014;1837,461–469

May AC, Aupperle RL, Stewart JL. Dark Times: The Role of Negative Reinforcement in Methamphetamine Addiction. Front Psychiatry. 2020;11:114.

McCoy MT, Jayanthi S, Wulu JA, Beauvais G, Ladenheim B, Martin TA, Krasnova IN, Hodges AB, Cadet JL. Chronic methamphetamine exposure suppresses the striatal expression of members of multiple families of immediate early genes (IEGs) in the rat: normalization by an acute methamphetamine injection. Psychopharmacology (Berl). 2011;215(2):353–65.

Meredith CW, Jaffe C, Ang-Lee K, Saxon AJ. Implications of chronic methamphetamine use: a literature review. Harv Rev Psychiatry. 2005;13:141–154.

Mitchell P. Coupling of phosphorylation to electron and hydrogen transfer by a chemi-osmotic type of mechanism. Nature. 1961;191:144–8.

Morava E, Kozicz T. Mitochondria and the economy of stress (mal) adaptation. Neurosci Biobehav Rev. 2013;37:668–680.

Nicholls DG. Mitochondrial dysfunction and glutamate excitotoxicity studied in primary neuronal cultures. Curr Mol Med. 2004;4(2):149–77.

Pei L, Wallace DC. Mitochondrial etiology of neuropsychiatric disorders. Biol. Psychiatry. 2018;83:722–730

Saito A, Imaizumi K. Unfolded Protein Response-Dependent Communication and Contact among Endoplasmic Reticulum, Mitochondria, and Plasma Membrane. Int J Mol Sci. 2018;19(10):3215.

Samad N, Saleem A, Yasmin F, Shehzad MA. Quercetin protects against stress-induced anxiety- and depression-like behavior and improves memory in male mice. Physiol Res. 2018;67(5):795–808.

Shin EJ, Dang DK, Tran TV, Tran HQ, Jeong JH, Nah SY, Kim HC. Current understanding of methamphetamineassociated dopaminergic neurodegeneration and psychotoxic behaviors. Archives of Pharmacal Research, 2017;40(4):403–428.

Shirakawa H, Sakimoto S, Nakao K, Sugishita A, Konno M, Iida S, Kusano A, Hashimoto E, Nakagawa T, Kaneko S. Transient receptor potential canonical 3 (TRPC3) mediates thrombin-induced astrocyte activation and upregulates its own expression in cortical astrocytes. J Neurosci. 2010;30(39):13116–29.

Shoptaw SJ, Kao U, Ling W. Treatment for amphetamine psychosis. Cochrane Database of Systematic Reviews. 2009;2009(1):CD003026.

Smith MJ, Thirthalli J, Abdallah AB, Murray RM, Cottler LB. Prevalence of psychotic symptoms in substance users: a comparison across substances. Compr Psychiatry 2009;50:245–250.

Song X, Chen Z, Jia R, Cao M, Zou Y, Li L, Liang X, Yin L, He C, Yue G, Yin Z.. Transcriptomics and proteomic studies reveal acaricidal mechanism of octadecanoic acid-3, 4 - tetrahydrofuran diester against Sarcoptes scabiei var. cuniculi. Scientific reports, 2017;7:45479.

Srisurapanont M, Ali R, Marsden J, Sunga A, Wada K, Monteiro M. Psychotic symptoms in methamphetamine psychotic in-patients. Int J Neuropsychopharmaco 2003;6:347–352.

Su H, Zhang J, Ren W, Xie Y, Tao J, Zhang X, et al. Anxiety level and correlates in methamphetamine-dependent patients during acute withdrawal. Medicine. 2017; 96:e6434.

Sun J, Chen F, Chen C, Zhang Z, Zhang Z, Tian W, Yu J, Wang K. Intestinal mRNA expression profile and bioinformatics analysis in a methamphetamine-induced mouse model of inflammatory bowel disease. Ann Transl Med 2020;8(24):1669.

Tian H, Li X, Tang Q, Zhang W, Li Q, Sun X, Zhao R, Ma C, Liu H, Gao Y, Han F. Yi-nao-jie-yu Prescription Exerts a Positive Effect on Neurogenesis by Regulating Notch Signals in the Hippocampus of Post-stroke Depression Rats. Frontiers in psychiatry, 2018;9:483.

Uhlmann A, Fouche JP, Koen N, Meintjes EM, Wilson D, Stein DJ. Fronto-temporal alterations and affect regulation in methamphetamine dependence with and without a history of psychosis. Psychiatry Research: Neuroimaging. 2016;248:30–38.

Valian N, Heravi M, Ahmadiani A, Dargahi L. Effect of methamphetamine on rat primary midbrain cells; mitochondrial biogenesis as a compensatory response. Neuroscience. 2019;406:278–289. Erratum in: Neuroscience. 2019;413:317–318.

Wallace DC. A mitochondrial etiology of neuropsychiatric disorders. JAMA Psychiatry 2017;74:863–864

Wearne TA, Cornish JL. A comparison of methamphetamine-induced psychosis and schizophrenia: a review of positive, negative, and cognitive symptomatology. Front Psychiatry. 2018;9:491.

Xie N, Yuan K, Zhou L, Wang K, Chen HN, Lei Y, Lan J, Pu Q, Gao W, Zhang L, Shen G, Li Q, Xiao H, Tang H, Xiang R, He M, Feng P, Nice EC, Wei Y, Zhang H, Yang J, Huang C. PRKAA/AMPK restricts HBV replication through promotion of autophagic degradation. Autophagy. 2016;12(9):1507–1520.

Zhang P, Kishimoto Y, Grammatikakis I, Gottimukkala K, Cutler RG, Zhang S, Abdelmohsen K, Bohr VA, Misra Sen J, Gorospe M, Mattson MP. Senolytic therapy alleviates Aβ-associated oligodendrocyte progenitor cell senescence and cognitive deficits in an Alzheimer’s disease model. Nat Neurosci. 2019;22(5):719–728.

Zhang Z, Yu J. NR4A1 Promotes Cerebral Ischemia Reperfusion Injury by Repressing Mfn2-Mediated Mitophagy and Inactivating the MAPK-ERK-CREB Signaling Pathway. Neurochem Res. 2018;43(10):1963–1977.

Zhou Y, Zhu H, Liu Z, Chen X, Su X, Ma C, Tian Z, Huang B, Yan E, Liu X, Ma L. A ventral CA1 to nucleus accumbens core engram circuit mediates conditioned place preference for cocaine. Nat Neurosci. 2019;22(12):1986–1999.

