## Supplementary Table 1 for "Quercetin ameliorates mitochondrial dysfunction and mitigates methamphetamine-induced anxiety-like behavior"

| **Supplementary Table 1. Details of DEGs in HIPP from Control vs Meth-treated mice** | | |
| --- | --- | --- |
| **Gene_id** | **fold change** | **p value** |
| Arc | 2.12697622 | 0.000598377 |
| Dusp1 | 1.280612858 | 0.001149044 |
| Egr1 | 1.778841297 | 0.0002803 |
| Egr2 | 1.68587008 | 0.024758373 |
| Egr4 | 1.430093708 | 0.015861385 |
| Fos | 2.486912126 | 0.000603645 |
| Fosb | 1.225353639 | 0.000438385 |
| Gadd45b | 1.067595202 | 0.001361706 |
| Hsp25-ps1 | 1.22755558 | 0.006954227 |
| Hspb1 | 1.218638077 | 0.006357643 |
| Junb | 1.419722905 | 0.012741178 |
| Nr4a1 | 1.525821037 | 0.005249888 |
| Rnu5g | 1.45598275 | 0.015010672 |
| Sdf2l1 | 1.098099472 | 0.013436189 |
