## Supplementary Table 2 for "Quercetin ameliorates mitochondrial dysfunction and mitigates methamphetamine-induced anxiety-like behavior"

| **Supplementary Table 2. Primer sequences** | | |
| --- | --- | --- |
| **Gene symbol** | **Primer sequence (Forward)** | **Primer sequence (Reverse)** |
| *c-fos* | ACAGCCTTTCCTACTACCA | GCACTAGAGACGGACAGA |
| *Arc* | TAAGTGCCGAGCTGAGAT | GTAGCCGTCCAAGTTGTT |
| *Nr4a1* | TCAATATGGAACACCAGCAA | AGGAGGCAGAGGAACAAG |
| *Egr1* | CAGCGCCTTCAATCCTCAAG | GCGATGTCAGAAAAGGACTCTGT |
| *Mfn2* | AGAACTGGACCCGGTTACCA | CACTTCGCTGATACCCCTGA |
| *Opa1* | TGGAAAATGGTTCGAGAGTCAG | CATTCCGTCTCTAGGTTAAAGCG |
| *Drp1* | CAGGAATTGTTACGGTTCCCTAA | CCTGAATTAACTTGTCCCGTGA |
| *IL-1β* | CCTTGTCGAGAATGGGCAGT | TTCTGTCGACAATGCTGCCT |
| *TNFα* | AACACACGAGACGCTGAAGT | TCCAGTGAGTTCCGAAAGCC |
| *IL-6* | CGGAGAGGAGACTTCACAGAG | CATTTCCACGATTTCCCAGA |
| *GAPDH* | AATGGATTTGGACGCATTGGT | TTTGCACTGGTACGTGTTGAT |
