## Supplementary Figures for "Quercetin ameliorates mitochondrial dysfunction and mitigates methamphetamine-induced anxiety-like behavior"

*Corresponding authors:

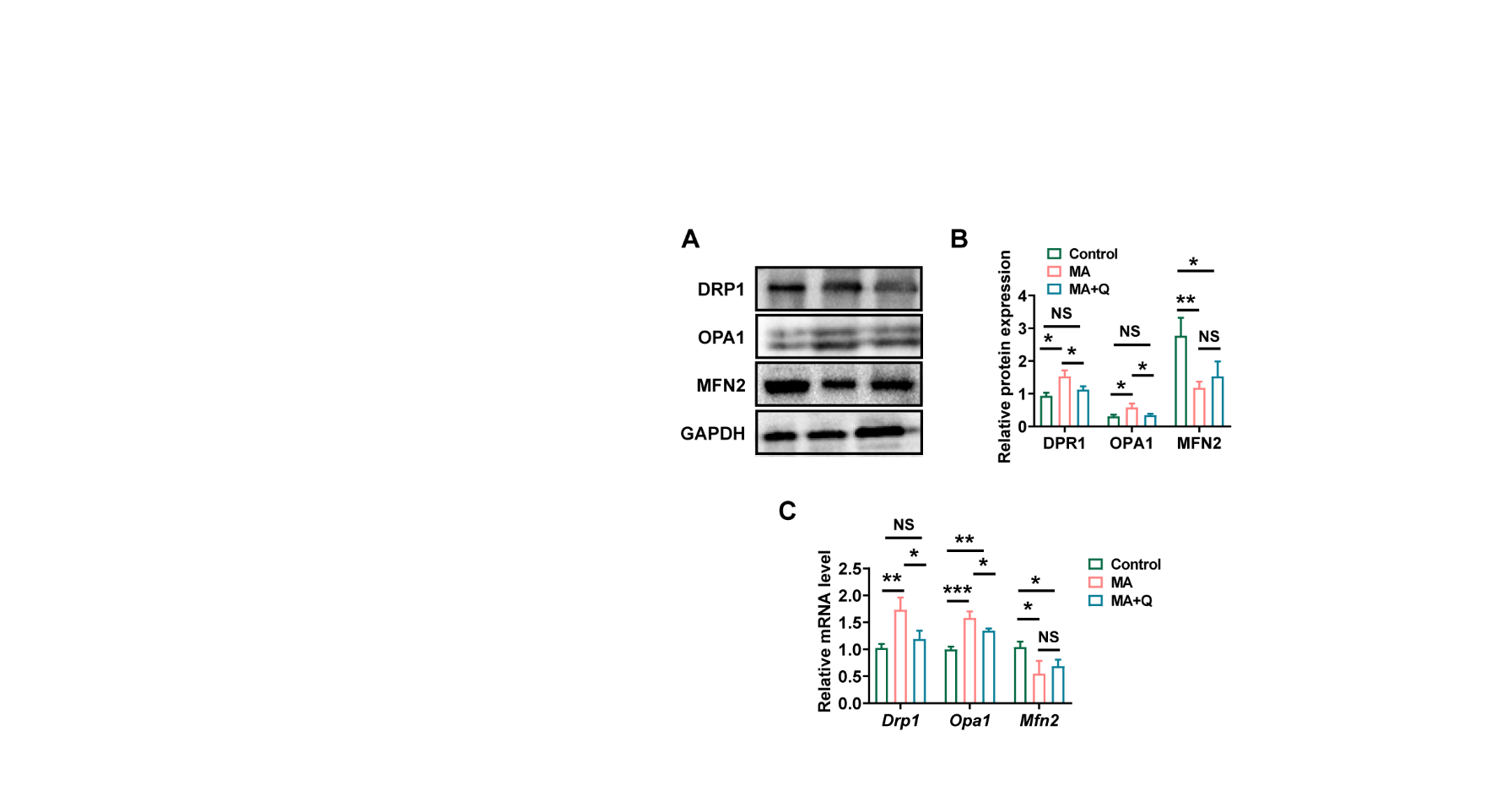

**Supplementary S1. Effect of MA and quercetin on mitochondria-associated genes in astrocytes.** A. Protein expression of mitochondria-associated genes, normalized to GAPDH protein expression. The quantitative expression values are presented in the bar graph (B). C. Relative expression of *Drp1* mRNA, *Opa1* mRNA, and *Mfn2* mRNA.

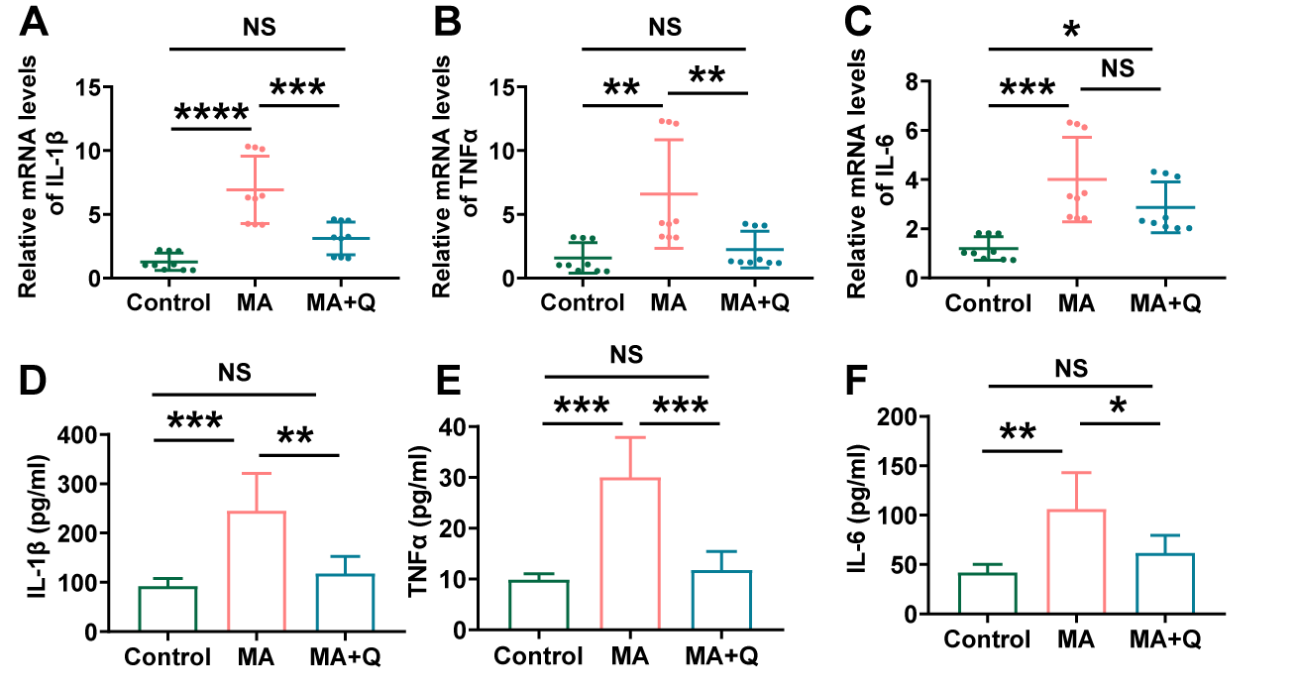

**Supplementary S2. Effect of MA and quercetin on pro-inflammatory cytokines in the hippocampus.** A-C. Expression of proinflammatory factors by qPCR in the hippocampus (*IL-1β*, *TNFα*, *IL-6*). D-F. The levels of the proinflammatory factors IL-1β were detected by ELISA.
